# A heterogeneously expressed gene family modulates biofilm architecture and hypoxic growth of *Aspergillus fumigatus*

**DOI:** 10.1101/2020.12.23.424276

**Authors:** Caitlin H. Kowalski, Kaesi A. Morelli, Jason E. Stajich, Carey D. Nadell, Robert A. Cramer

## Abstract

The genus *Aspergillus* encompasses human pathogens such as *Aspergillus fumigatus* and industrial powerhouses such as *Aspergillus niger.* In both cases, *Aspergillus* biofilms have consequences for infection outcomes and yields of economically important products. Yet, the molecular components influencing filamentous fungal biofilm development, structure, and function remain ill-defined. Macroscopic colony morphology is an indicator of underlying biofilm architecture and fungal physiology. A hypoxia-locked colony morphotype of *A. fumigatus* has abundant colony furrows that coincide with a reduction in vertically-oriented hyphae within biofilms and increased low oxygen growth and virulence. Investigation of this morphotype has led to the identification of the causative gene, *biofilm architecture factor A (bafA),* a small cryptic open reading frame within a subtelomeric gene cluster. BafA is sufficient to induce the hypoxia-locked colony morphology and biofilm architecture in *A. fumigatus.* Analysis across a large population of *A. fumigatus* isolates identified a larger family of *baf* genes, all of which have the capacity to modulate hyphal architecture, biofilm development, and hypoxic growth. Furthermore, introduction of *A. fumigatus bafA* into *A. niger* is sufficient to generate the hypoxia-locked colony morphology, biofilm architecture, and increased hypoxic growth. Together these data indicate the potential broad impacts of this previously uncharacterized family of small genes to modulate biofilm architecture and function in clinical and industrial settings.

**Importance:** The manipulation of microbial biofilms in industrial and clinical applications remains a difficult task. The problem is particularly acute with regard to filamentous fungal biofilms for which molecular mechanisms of biofilm formation, maintenance, and function are only just being elucidated. Here we describe a family of small genes heterogeneously expressed across *Aspergillus fumigatus* strains that are capable of modifying colony biofilm morphology and microscopic hyphal architecture. Specifically, these genes are implicated in the formation of a hypoxia-locked colony morphotype that is associated with increased virulence of *A.* f*umigatus*. Synthetic introduction of these gene family members, here referred to as biofilm architecture factors, in both *A. fumigatus* and *A. niger* additionally modulates low oxygen growth and surface adherence. Thus, these genes are candidates for genetic manipulation of biofilm development in Aspergilli.

## Introduction

Biofilms are surface-adhered populations or communities of microorganisms that are embedded in an extracellular matrix, have unique transcriptional programs, and are typically tolerant to exogenous stress (1–3). Bacterial biofilms have received the majority of attention over the past decades with a focus on how bacteria initiate biofilm growth (4, 5), exogenous factors that influence biofilm development (6, 7), and methods of sensitizing biofilms to exogenous stressors (8, 9). Filamentous fungal biofilm research is still in its relative infancy compared to bacterial biofilms, with the majority of research focusing on the yeast and polymorphic fungi (10–13). Filamentous fungi, or molds, form biofilms, and this mode of growth is important for clinical and industrial applications (1, 14–16). The *Aspergillus* genus of filamentous fungi includes human pathogens, *Aspergillus fumigatus*, and biotechnological powerhouses, *Aspergillus niger* and *Aspergillus oryzae.* In regard to the former, during life-threatening infections *A. fumigatus* biofilms form within the airways and lung tissue during aspergilloma and invasive aspergillosis, respectively (17, 18). In industry, biofilm formation is a proverbial doubleedged sword, where surface-immobilized *A. niger* biofilms produce higher yields of citric acid than free-floating planktonic cultures (19), but recalcitrant biofilms can be difficult to remove and corrosive (20). Despite the significance of filamentous fungal biofilms, and specifically those of *Aspergillus* biofilms, large gaps in knowledge remain regarding the molecular components influencing filamentous fungal biofilm formation, structure, and function.

A colony of a single microbial species cultured on a semi-sold surface can be considered a biofilm, and changes in colony morphology predict or reflect important biofilm characteristics and organism physiology (21, 22). Diverse exogenous factors have been described that influence microbial colony morphotypes. For fungi, these include zinc induction of radial colony grooves during the filamentous growth of the basidiomycete *Tricholoma matsuke* (23), low oxygen, or hypoxic, induction of colony wrinkling in the polymorphic yeast *Candida albicans* (24, 25) and colony furrowing in the filamentous fungus *A. fumigatus* (26). The pool of molecular regulators of the wrinkled colony morphotype of *C. albicans* have been define and linked to increased oxygen penetration and virulence (24, 27). Despite numerous reports of similarly complex colony morphotypes among filamentous fungi (26, 28), it remains a significant gap in knowledge how these morphotypes reflect the physiology of the population and importantly what molecular mechanisms contribute to their development.

Previously, we have demonstrated that oxygen tensions significantly contribute to colony morphology features in *A. fumigatus* (26). In an experimentally evolved strain of *A. fumigatus,* EVOL20, that was serially passaged in hypoxic conditions, a colony morphotype was formed in normal oxygen that shared features of a typical hypoxia-grown colony. These colony features consistent with a hypoxia-grown colony include increased colony furrows and a white perimeter of vegetative growth (Fig. S1A). We defined this hypoxia locked morphotype as H-MORPH and the parental or normal oxygen morphotype as N-MORPH (26). EVOL20 and other H-MORPH strains coincidently have altered hyphal arrangements within submerged biofilms characterized by a reduction in vertically-oriented hyphae. We identified a putative transcriptional regulator, *hrmA*, of a subtelomeric gene cluster (*hrmA-*associated gene cluster (HAC)) that is required for H-MORPH in EVOL20 (Fig. S1B). In addition to H-MORPH, *hrmA* expression coincides with increased hypoxia fitness and reduced adherence relative to the parental strain AF293 (Fig. S1C) (26). A collagen-like protein encoding gene *(cgnA)* located within HAC appeared to be essential for H-MORPH in EVOL20, as targeted deletion of the annotated *cgnA* coding sequence reverted the H-MORPH of EVOL20 to N-MORPH (Fig. S1C) (26). However, constitutive expression of *cgnA* in the N-MORPH AF293 did not result in an H-MORPH phenotype. Thus, it remained unclear how *hrmA* and *cgnA*, and potentially other HAC genes, brought about H-MORPH and the associated phenotypes of EVOL20. Here we describe a continuation of this work, in which we identified an unannotated, cryptic gene within HAC that is shared among putative HAC orthologous clusters in multiple *A. fumigatus* strains. As this cryptic gene is sufficient to generate H-MORPH in the parental strain AF293 we propose the name *biofilm architecture factor A* (*bafA*). *bafA* expression is sufficient to generate H-MORPH in a distant *Aspergillus* species, *A. niger*, demonstrating the potential for synthetic modulation of these genes to modify *Aspergillus* biofilms in both clinical and industrial settings.

## Material and Methods

### Strains, strain construction, and culture conditions

All strains used in this study are listed in Table S1. The *Aspergillus niger* strain A1144 was purchased from the Fungal Genomic Stock Center, Kansas State University, Manhattan, KS (29). All strains were maintained on glucose minimal media (GMM; 1% glucose, 6g/L NaNO3, 0.52g/L KCL, 0.52g/L MgSO4·7H2O, 1.52g/L KH2PO4 monobasic, 2.2mg/L ZnSO4·7H20, 1.1mg/L H3BO3, 0.5mg/L MnCl2·4H2O, 0.5mg/L FeSO4·7H2O, 0.16mg/L CoCl2·5H2O, 0.16mg/L CuSO4·5H2O, 0.11mg/L (NH4)6Mo7O24·4H2O, 5mg/L Na4EDTA, 1.5% agar; pH 6.5) and spores were collected and counted for experimentation in 0.01% Tween80. To generate the various overexpression strains, we started with the pTMH44.2 plasmid which includes *A. nidulans gpdA* promoter and terminator *trpC* separated by green fluorescent protein (GFP) fragment (30). We inserted *ptrA* for pyrithiamine resistance from pPTR I (Takara) or *hygB* for hygromycin resistance from pBC-hygro (Creative Biogene) 3’ to the *trpC* terminator to generate pTDS8 and pTDS9, respectively. The GFP fragment could then be replaced through restriction enzyme ligation with AscI (New England Biolabs) at the 5’ end and NotI (New England Biolabs) at the 3’ end. We amplified the *bafA* sequence from AF293 genomic DNA and *bafB* and *bafC* sequences from CEA10 genomic DNA with primers that introduced AscI/NotI sites. Overlap PCR was used to generate the *bafB-GFP* fragment with AscI/NotI sites. For the generation of the *cgnA^RECON^* strain, pBluescript II KS(+) (Addgene) was utilized. Briefly, we expanded the multiple cloning site and introduced the hydromycin resistance cassette amplified with XhoI/XbaI restriction sites from pBC-Hygro (Creative Biogene). From AF293 genomic DNA we amplified ~1kb 5’ and ~500 bp 3’ of *cgnA* to generate the *cgnA* reconstitution cassette with AscI and PacI.

Digested amplification products and digested vectors (pTDS8 or pTDS9) were ligated with T4 DNA ligase (New England Biolabs) and transformed into CaCl2 competent DH5a *E. coli.* Plasmids were confirmed by restriction digestion and Sanger sequencing. Plasmids were isolated (Zyppy Plasmid Miniprep, Zymo Research) and ectopically into the fungal genome using previous protocols for the generation and transformation of protoplasts using Lysing Enzyme from *Trichoderma harzianum* (Sigma: L1412) for *A. fumigatus* germlings (31) and Vinotaste Pro (Gusmer Enterprises: No. VINOTASTEPRO-250) for *A. niger* hyphae (32). Primers used for the construction of strains are provided in Table S2.

### Oxygen measurements of colony biofilms

A Unisense Oxygen Measuring System 1-CH (Unisense no. OXY METER) equipped with a Micromanipulator (Unisense no. MM33), Motorized Micromanipulator Stage (Unisense no. MMS), Motor Controller (Unisense no. MC-232), and a 25 μm Clark Type/Amperometric Oxygen Sensor (Unisense no. OX-25) was used to quantify oxygen above and within colony biofilms. The readings were automated and analyzed through SensorTrace Suite Software v3.1.151 (Unisense no. STSUITE). Colony biofilms were point inoculated with 1000 spores in 0.002 mL on glucose minimal media with 1.5% agar and cultured in normal oxygen or hypoxia (0.2% O_2_) with 5% CO_2_ at 37° for 72 hours. The calibrated oxygen sensor was positioned 1.2 mm or 0.5 mm above the agar surface and measurements in technical duplicates were acquired every 0.1 mm over a period of 5 s with a 5 s wait period at each new depth. Colony biofilms were analyzed in a minimum of biological triplicates.

### ORF prediction and sequence alignments

RNA-sequencing reads from (26) for EVOL20 normoxia sample and AF293 normoxia sample were uploaded to Integrative Genome Viewer with the annotated AF293 genome file. The absence of an annotated gene between *Afu5g14910* and *Afu5g14920* was confirmed using FungiDB.org (33). The sequence corresponding to the mapped reads in this region, with consideration for the intro-like space, were uploaded to NCBI ORFfinder using a minimal ORF length of 75 nucleotides, standard genetic code, and “ATG” only start codon. This provided the DNA and protein translation. The protein and DNA sequences provided by ORFfinder were BLAST against the *A. fumigatus* genome using FungiDB. Alignments were then generated between the BLAST hits on FungiDB and the cryptic gene query sequence in NBCI BLASTblastp suite. Given the high DNA and amino acid identity between the cryptic gene sequences (Afu5g14915, *bafA*) and *bafB* we defined the two exons and intron of *bafA* from the *bafB* sequence.

### Colony Morphology Assays and Quantification

Glucose minimal media agar plates (1.5% agar) were spot inoculated at the center of the plate with 1000 spores in 0.002 mL of 0.01% Tween 80. Plates were incubated at 37° in the dark for 72-96 hours at 21% or 0.2% O_2_ with 5% CO_2_. Images were captured with a Canon PowerShot SX40 HS. Images are representative of three independent biological samples. Images were converted to 8-bit in Fiji (ImageJ). Quantification of colony furrows and calculation of the percent vegetative mycelia were quantified as previously described using Fiji (26).

### Liquid Morphology

Aliquots of 10 mL were taken and photographed from 18 hour cultures of 10^6^ spores grown in 50 mL LGMM in normal oxygen conditions at 37° with constant agitation at 200 rpm.

### Hypoxia Growth Assays

Hypoxia growth assays to calculate the ratio of hypoxia to normoxia growth (hypoxia fitness, H/N) of a strain were performed in 100 mL of glucose minimal media in acid-washed baffled glass flasks with a total of 5×10^6^ spores per mL. Cultures were incubated at 37° in the dark at 21% or 0.2% O_2_ with 5% CO_2_ and shaking at 200 rpm. Incubation for *A. fumigatus* strains was 48 hours in both 21% or 0.2% O_2_ and for *A. niger* strains was 72 hours in both 21% or 0.2% O_2_. Fungal mycelia were collected through Miracloth, frozen at −80° and lyophilized for 16 hours before being weighed.

### Adherence Assays

Adherence was measured using a crystal violet assay as previously described (34). Briefly, 10^4^ spores were inoculated in 0.1 mL of liquid glucose minimal media per well of a U-Bottom 96-well plate, centrifuged at 250 x g for 10 minutes, and then incubated at 37° and 5% CO_2_ in the dark for 24 hours. The wells were washed twice with water, stained for 10 minutes with 0.1% (wt/vol) crystal violet, washed twice more with water and then de-stained with 100% ethanol. Optical density was measured at 600 nm.

### RNA extraction and Gene Expression Assays

RNA was extracted from mycelia grown in static biofilm cultures in 15 mL of liquid glucose minimal media in a 100 mm plastic petri dish with 10^5^ spores per mL. Mycelium was collected (~50 mg) and flash frozen in liquid nitrogen. Samples were transferred to – 80° for at least 1 hour before bead beating with 2.3 mm zirconia-silica beads (Biospec: No. 11079125z) for 1 minute in 0.2 mL TRIsure (Bioline: BIO-38033). Homogenized tissue was brought to 1 mL volume with 0.8 mL TRIsure. Chloroform (0.2 mL) was added to the TRIsure tissue homogenate and centrifuged for 15 minutes at 21130 rcf at 4°. The aqueous phase was transferred to 0.6 mL 2-Propanol and centrifuged for 10 minutes at 21130 rcf at 4°. The RNA pellet was washed with 0.5 mL 75% ethanol and resuspended in RNAse-free water. 5 μg of RNA was DNAse treated with Ambion TURBO DNA-free™ Kit (Invitrogen: AM1907) following manufacturer’s instructions. cDNA synthesis was carried out using the QuantiTect. Reverse Transcription kit (Qiagen: No. 205311) with 500 ng of RNA. Gene expression was quantified using IG^TM^ SYBR^®^ Green Supermix (Bio-Rad: 1708880) with a CFX Connect Real-Time PCR Detection System (Bio-Rad) equipped with CFX Maestro Software (Bio-Rad). Reactions were 0.02 mL volume and contained 25 μg of cDNA. The mRNA levels were normalized to *actA* and *tub2* for *A. fumigatus* and to *tubB* for *A. niger.* Normalized expression was quantified as previously described (35).

### Biofilm Microscopy and Architectural Analysis

Fluorescence confocal microscopy was performed on an Andor W1 Spinning Disk Confocal with a Nikon Eclipse Ti inverted microscope equipped with a CFI Plan Fluor 20XC MI objective (Nikon). *A. fumigatus* and *A. niger* biofilms were cultured for imaging at 10^5^ spores per mL in MatTek dishes (MatTek: No. P35G-1.0-14-C) in 2 mL liquid glucose minimal media for 24 hours at 37° with 5% CO_2_ in the dark. For visualization at 405 nm, 15 minute prior to imaging biofilms were stained with 25μg/mL calcofluor white (Fluorescent Brightener 28) (Sigma: No. F3543). The CFI Plan Fluor 20XC MI objective was used with water to image the bottom ~300 μm of the biofilm with Z-slices collected every 1.2-1.5 μm. 3D rendering and image processing were performed in Nikon Elements Viewer (Nikon). Quantification of biofilm architecture and generation of the heat map figures was carried out as previously described using the BiofilmQ framework written in MatLab (36). The framework is freely available for download at www.drescherlab.org/data. All biofilm images and single columns in heat maps are representative of a minimum of three independent biological replicates.

### Protein localization

Spores were cultured at 10^4^ spores/mL in 0.2 mL liquid glucose minimal media for 10 hours at 37° with 5% CO_2_ in the dark on MatTek dishes (MatTek: No. P35G-1.0-14-C). At this point, hyphae of various sizes had formed. Hyphae were imaged unfixed on an Andor W1 Spinning Disk Confocal with a Nikon Eclipse Ti inverted microscope equipped with a CFI Plan Fluor 100X Oil objective (Nikon). 3D rendering and image analysis were performed in Nikon Elements Viewer (Nikon). For FM4-64 staining, hyphae were incubated in 10um FM4-64 in PBS for 15 minutes on ice and then briefly incubated in 37° for 5 minutes before being washed 2x with PBS and imaged as described above. For detection of B-glucan through Dectin-1 binding protocols were follow as previously published (37). Briefly, 12-hour hyphal cultures were blocked in FACS buffer containing fetal bovine serum for 30 minutes, washed 2x with PBS, and then incubated in 150 uL of soluble dectin-1 for 1 hr at room temperature. Hyphae were stained with Goat antiHuman IgG Alexa Fluor 594 in PBS for 1 hour at room temperature.

### Quantification of conidiation

Conidiation was quantified by spread plating 0.3 mL of spores at 10^6^ spores/mL onto a glucose minimal media agar plate. Plates were incubated at 37° in the dark at 21% O_2_ or 0.2% O_2_ with 5% CO_2_ for 48 or 72 hours. Spores were collected from each plate in 5 mL of 0.01% Tween 80, and 0.1mL of the spores, or if needed a 1:10 dilution in 0.01% Tween 80, were transferred to a flat bottom 96-well plate. Spores were using forward and side scatter on a MacsQuant VYB Flow Cytometer with a slow flow rate and gentle mixing. Gating was set for single, non-swollen spores and analyzed in FlowJo v9.9.6. Three independent biological samples were counted for each strain in technical triplicates.

### Strain genome assembly, baf presence, and phylogenetic tree construction

Unassembled sequence reads from public NCBI Sequence Read Archive (SRA) and strain datasets generated in the Cramer and Stajich labs were processed to produce draft assembled genomes as part of ongoing research in A. fumigatus evolution (https://github.com/stajichlab/Afum_popgenome). The pipeline utilizes the AAFTF v0.2.3 (Automatic Assembly For The Fungi) pipeline https://github.com/stajichlab/AAFTF (Stajich JE and Palmer J. (2019, September 17). stajichlab/AAFTF: v0.2.3 release. doi: 10.5281/zenodo.3437300) which trims sequences for quality, filters for phiX and vector contamination, and assembles genomes with SPAdes v 3.13.1 followed by trimming of adapter and contamination sequences. The assembly is further removed of redundancy and polished with Pilon (38). These assembled genomes were searched for copies of baf including cryptic loci identified through translated searches.

The evolutionary relationship of the 92 strains was inferred by constructing a phylogenetic tree of the genomic variants. The complete set of public *A. fumigatus* strains were initially used but pruned from the final tree after removing nearly identical isolates based on visual inspection of the phylogenetic tree. The variants were identified by downloading Illumina sequence data from NCBI Sequence Read Archive (Table SX) and aligning these to the reference *A. fumigatus* strain Af293 genome downloaded from FungiDB (Release 39) (33). Variants were identified by aligning reads to the genome with bwa v0.7.17 (39) followed by conversion to BAM and CRAM file format after running the fixmate and sort steps with samtools v1.10 (40). The alignments were filtered by identifying and removing duplicate reads using MarkDuplicates tool in the picard tools v2.18.3 (http://broadinstitute.github.io/picard). Reads were also realigned around gaps using RealignerTargetCreator and IndelRealigner in the Genome Analysis Toolkit GATK v3.7 (41). Were genotyped relative to the A. fumigatus reference genome AfF293 using HaplotypeCaller on individual CRAM files followed by jointly calling variants with the GenotypeGVCFs in GATK v4.0 (doi: 10.1101/201178). Identified variants were filtered using GATK’s SelectVariants to create a Variant Call File Format (VCF) file split into one for Single Nucleotide Polymorphisms (SNP) and Insertion/Deletions (INDEL) with the following parameters: for SNPs: -window-size = 10, -QualByDept < 2.0, -MapQual < 40.0, -QScore < 100, -MapQualityRankSum < −12.5, -StrandOddsRatio > 3.0, – FisherStrandBias > 60.0, -ReadPosRankSum < −8.0. For INDELs: -window-size = 10, – QualByDepth < 2.0, -MapQualityRankSum < −12.5, -StrandOddsRatio > 4.0, – FisherStrandBias > 200.0, -ReadPosRank < −20.0, -InbreedingCoeff < −0.8. The filtered SNP report was processed with bcftools v1.11 (http://www.htslib.org/) to generate an alignment of the strains. A total of 71,513 parsimony-informative and 268 singleton sites were in the alignment across the 92 strains including the Af293 reference genome, A Maximum Likelihood phylogenetic tree was constructed from this alignment with IQ-TREE v 2.1.1 using the GTR+ASC model and 1000 bootstrap (-m GTF+ASC −B 1000) (42).

To identify copies of the homologs in the assembled genomes of strains, DNA sequences of *bafB* and *bafC* genes were searched against the compiled dataset of *A. fumigatus* genomes with the following procedures which are part of github project (https://github.com/stajichlab/Afum_baf; doi: 10.5281/zenodo.3726371). The pipeline.sh file includes the analysis steps and baf_mRNA.fa are the query sequences including the founder copies AFUB_044360_bafB and AFUB_096610_bafC. Briefly this includes a nucleotide search of the defined loci with FASTA against the assemblies to identify and then extract the sequences with a custom Perl script (43). The results are combined, assigned a name based on best hit search to the starting database of named *baf* sequences. A multiple alignment was generated using MAFFT (https://mafft.cbrc.jp/alignment/software/). A phylogenetic tree of the gene sequences was constructed with FastTree v2.1.11 (44).

### Statistical Analysis

All statistical analyses were performed in GraphPad Prism 8 unless otherwise noted. Error bars indicate standard error around the mean and individual data points indicate independent biological samples when shown. Images of colony biofilms or submerged biofilms are representative of a minimum of three independent biological samples.

### Data Availability

The primary sequence FASTQ sequence reads for the strains are all available in the NCBI SRA database and detailed in Table S2.

## Results

### Colony furrows increase oxygen diffusion within colony biofilms

A definitive feature of *A. fumigatus* H-MORPH and a common feature of hypoxia-grown colony morphotypes is the presence of furrows or invaginations within the colony biofilms (Fig. 1A, Fig. S1A). We hypothesize that these furrows increase colony surface area and oxygen diffusion into the colony. To test this hypothesis, a microelectrode oxygen sensor was utilized to quantify oxygen above, within, and below the colony biofilms of the reference N-MORPH strain AF293 grown in normoxia (normal oxygen, 21% O_2_, non-furrowing condition) or hypoxia (furrowing condition, 0.2% O_2_) (Fig. 1B). The non-furrowing normoxia-grown colony of AF293 shows a precipitous drop in oxygen within the 400 μm of the colony above the agar surface (0 μm) and 200 μM below the agar surface (embedded colony). In contrast, the furrowed, hypoxia-grown colonies of AF293, measured both within the furrowing (F) or non-furrowing (NF) regions, show significantly increased oxygen levels within the colonies (Fig. 1A, B). To determine if the furrows in the normoxia-grown H-MORPH colony of EVOL20 also impact oxygen diffusion, we utilized the same approach to quantify oxygen in AF293 normoxia-grown colonies and furrowing (F) and non-furrowing (NF) regions of EVOL20 normoxia-grown colonies (Fig 1A, C). Within the furrows of the EVOL20 colony oxygen is significantly increased above, at, and below the agar surface (0 μm). Additionally, the non-furrowing regions of the EVOL20 colony biofilm also have significantly increased oxygen compared to AF293 within the embedded colony (0-200 μm) (Fig. 1C). Together these data suggest that colony furrowing of *A. fumigatus* occurs in hypoxia in part to increase oxygen diffusion into the colonies, and that the furrows of the hypoxia-evolved H-MORPH strain EVOL20 develop even under normoxia to increase oxygen deep within the colonies. The increased oxygen diffusion within H-MORPH colonies coincides with altered hyphal architecture within biofilms, increased hypoxic growth, reduced adherence, and increased inflammation and virulence (Fig. S1C) (26).

**Figure 1.**
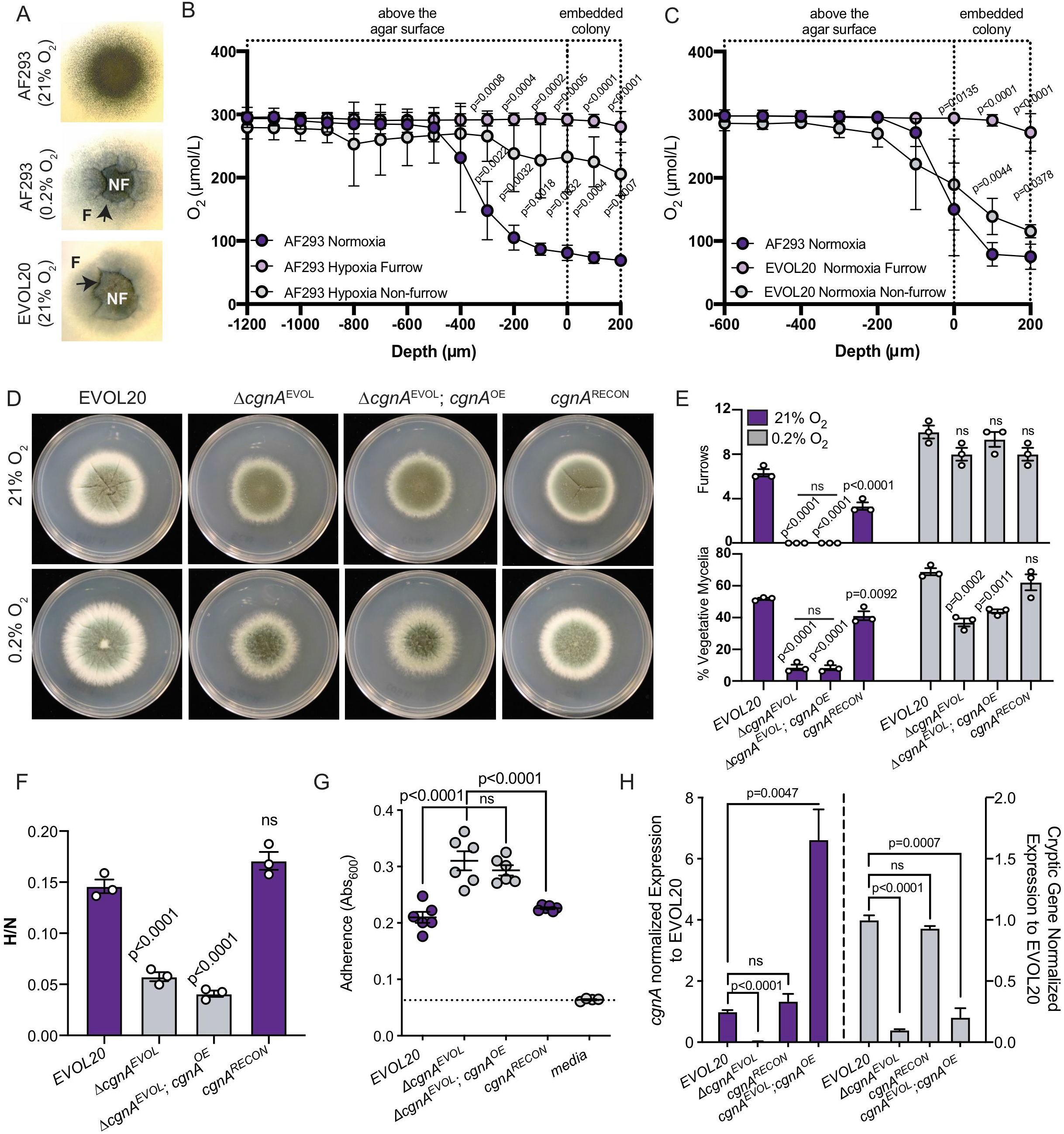
A cryptic gene within the *hrmA-associated* gene cluster is necessary for the hypoxia-evolved phenotypes of EVOL20. (A) 72-hour colony biofilms used for oxygen measurements with furrows (F) and non-furrowing (NF) regions labeled. Images are representative of three independent biological samples. (B) Oxygen quantification of AF293 colony biofilms grown in normoxia (21% O_2_) or hypoxia (0.2% O_2_). In hypoxia-grown colonies oxygen was measured both in furrows (F) and non-furrows (NF). n=3 independent biological replicates. Error bars indicate standard error around the mean. Multiple One-way ANOVAs performed with Dunnett’s posttest at each depth. (C) Oxygen quantification for normoxia-grown colonies of AF293 and EVOL20. EVOL20 colonies were measured in furrowing (F) and non-furrowing (NF) regions. N=3 independent biological replicates. Error bars indicate standard error around the mean. Multiple Oneway ANOVAs performed with Dunnett’s posttest at each depth. (D) 96-hour colony biofilms in normoxia (21% O_2_) and hypoxia (0.2% O_2_). Images are representative of three independent biological samples. (E) Quantification of colony biofilm morphological features from three independent biological samples. One-way ANOVA with Dunnett’s posttest for multiple comparisons was performed relative to EVOL20 within each oxygen environment. (F) The ratio of fungal biomass in hypoxia (0.2% O_2_) relative to fungal biomass in normoxia (21% O_2_) (H/N) in shaking flask cultures. One-way ANOVA with Dunnett’s posttest for multiple comparisons was performed relative to EVOL20. n=3 independent biological samples. (G) Adherence to plastic measured through a crystal violet assay. Dashed line marks the mean value for media alone. One-way ANOVA with Dunnett’s posttest for multiple comparisons was performed relative to Δ*cgnA^EVOL^.* n=6 independent biological replicates. (H) Gene expression measured by qRT-PCR for *cgnA* and the cryptic ORF. n=3 independent biological replicates. One-way ANOVA with Tukey’s multiple comparison test was performed.

### The native 5’ sequence to cgnA is required to complement the loss of cgnA in EVOL20

We have previously characterized a role for the HAC gene cluster in the generation of H-MORPH in EVOL20, based on the observation that *hrmA,* the HAC regulator, and the annotated *cgnA* coding sequene are required for H-MORPH in EVOL20. However, while constitutive expression of *hrmA* in the reference strain AF293 is sufficient to elevate mRNA levels of the HAC genes and generate H-MORPH, the expression of *cgnA* alone is not sufficient to generate H-MORPH (26). Therefore, we hypothesized that elevated expression of multiple HAC genes may be required to generate the H-MORPH phenotype.

As the majority of annotated HAC genes, with the exception of Afu5g14920, remain unaltered following the loss of *cgnA* in EVOL20 (Δ*cgnA*^EVOL^) (26), we overexpressed *cgnA* in this background (Δ*cgnA*^EVOL^; *cgnA^OE^)* using an *Aspergillus nidulans gpdA* promoter to drive constitutive expression (Fig. 1D). We discovered that the overexpression of *cgnA* could not restore the H-MORPH phenotype in Δ*cgnA*^EVOL^. Instead, the Δ*cgnA*^EVOL^; *cgnA^OE^* strain colony morphology is not significantly different compared to Δ*cgnA*^EVOL^ respective to furrowing and percent vegetative mycelia (Fig. 1D, E). We next hypothesized that the native sequence 5’ of *cgnA* may be required to restore H-MORPH in Δ*cgnA*^EVOL^. Ectopic integration of *cgnA* with its native promoter and 5’ sequence *(cgnA^RECON^)* is able to reconstitute H-MORPH in Δ*cgnA*^EVOL^ with elevated colony furrows and an increased percentage of vegetative mycelia in normoxia (Fig. 1D, E). In addition to a transition from the H-MORPH phenotype of EVOL20 to N-MORPH, Δ*cgnA^EVOL^* has a significantly reduced ratio of hypoxic to normoxia growth (hypoxia fitness, H/N) (Fig. 1F) and significantly increased hyphal adherence (Fig. 1G) compared to EVOL20 (26, 33). Where Δ*cgnA^EVOL^; cgnA^OE^* does not restore either of these phenotypes to the level of EVOL20, *cgnA*^RECON^, where *cgnA* is reintroduced with its native 5’ sequence, restores both hypoxia fitness and adherence of Δ*cgnA^EVOL^* similarly to that of EVOL20 (Fig. 1F, G). While integration loci of *cgnA* in the Δ*cgnA^EVOL^; cgnA^OE^* and *cgnA^RECON^* strains may not be identical and contribute to some phenotypic variation, the absolute necessity of the native sequence 5’ of *cgnA* to complement the loss of *cgnA* in EVOL20 (Δ*cgnA^EVO^*^L^) prompted us to investigate this genomic region more closely.

### A cryptic gene is encoded 5’ of cgnA within HAC and is required for H-MORPH and HAC related phenotypes

Utilizing published RNA-sequencing data, we identified a substantial region of mapped reads 5’ to *cgnA* in EVOL20 that were absent in AF293 (Fig. S2A) (26). Neither the AF293 assembled reference genome, nor the partially assembled genome of A1163, annotate a gene within this region (26, 33). It is unlikely these reads belong to the same transcript as *cgnA* as they map to the opposite strand. Therefore, we hypothesize that these reads map to an independent cryptic gene within HAC, and that this gene may be important for H-MORPH and other EVOL20-related phenotypes (i.e. hypoxia fitness, adherence, and biofilm architecture) (26). To determine if our strategies to delete *cgnA* interrupted the mRNA levels of this cryptic gene, we designed primers within the predicted open reading frame (ORF) to quantify relative expression in two isogenic strain sets: EVOL20/Δ*cgnA^EVOL^* and in *hrmA^R-EV^/hrmA^R-EV^; Δ*cgnA (Table S1). In both cases deletion of the *cgnA* coding sequence reduces *cgnA* mRNA levels and mRNA levels corresponding to the cryptic gene (Fig. S2B, C).

With Integrative Genome Viewer and NCBI ORF Finder, we were able to predict a two exon ORF of 579 base pairs (bp) from the region corresponding to the cryptic gene (Fig. S2D). In the DNA construct used to generate Δ*cgnA*^EVOL^;*cgnA*^OE^, *cgnA* expression was driven by the constitutive *A. nidulans gpdA* promoter and therefore the native 5’ sequence containing the cryptic gene ORF was not re-introduced (Fig. S2E). In contrast, the DNA construct used to generate the *cgnA*^RECON^ strain utilized the native sequence 5’ to *cgnA* to drive expression. This region includes the entire predicted coding sequence of the cryptic gene (Fig. S2E). Gene expression analysis confirmed that both Δ*cgnA*^EVOL^;*cgnA*^OE^ and *cgnA*^RECON^ have *cgnA* mRNA levels equivalent to or greater than those of EVOL20, but only *cgnA*^RECON^ restores mRNA levels of the cryptic gene similarly to EVOL20 (Fig. 1H). Only with the strain *cgnA*^RECON^, where both *cgnA* and the cryptic gene are expressed, is H-MORPH restored (Fig. 1D, E), hypoxic fitness increased (Fig. 1F), and adherence reduced (Fig. 1G) in Δ*cgnA*^EVOL^ to resemble EVOL20. Thus, the *cgnA* sequence alone is not sufficient to generate the EVOL20 phenotypes but requires the 5’ cryptic gene. Based on the previously published phenotypes of EVOL20 and the data presented herein, we propose the name biofilm architecture factor (*bafA*) for this cryptic gene.

### The HAC cryptic gene shares significant similarity to genes encoded within putative clusters orthologous to HAC in an independent strain

Previous phylogenetic analysis of the HAC genes *hrmA* and *cgnA* for presence across *A. fumigatus* strains revealed heterogeneity across the species, where some strains did not encode *hrmA* or *cgnA* (26). However, some *A. fumigatus* strains, such as the well-studied CEA10, encoded putative orthologs to *hrmA* within putative orthologous HAC clusters where the neighboring genes encode proteins with domain and amino acid sequence similarities to the proteins encoded by the genes of HAC (26). This observation suggests that some strains, like CEA10, may encode multiple HAC-like clusters in addition to HAC. While these orthologous HAC-like clusters show no evidence of encoding a putative ortholog of *cgnA*, they encode a gene that is similar to the HAC cryptic gene *bafA* (Fig. S3A). In the *hrmB* associated cluster (HBAC) from CEA10, the predicted amino acid sequence of BafA in AF293 shares 78.35% identity with that of AFUB_044360 (Fig. S3B). Additionally, in the *hrmC* associated cluster (HCAC) from CEA10, the predicted amino acid sequence of BafA in AF293 shares 45.41% identity with that of AFUB_096610 (Fig. S3C). At the level of DNA sequence, the similarity is even greater with 87% nucleotide identity between *bafA* and AFUB_044360 and 68% nucleotide identity between *bafA* and AFUB_096610. Based on these sequence similarities we propose the name *bafB* for AFUB_044360 and *bafC* for AFUB_096610. Notably, all three of these genes, *bafA, bafB,* and *bafC,* are located adjacent to a gene encoding a hypothetical protein with a conserved domain of unknown function DUF2841. The role of these DUF2841-containing proteins and their potential role in the development of H-MORPH remains the focus of future study.

AF293 and EVOL20 only encode HAC and there is no evidence for intact HBAC or H_C_AC in these genomes based on orthologs to *hrmA.* In contrast, CEA10 encodes all three putative clusters. Previously we identified H-MORPH strains of *A. fumigatus* that do not encode *hrmA*/HAC, and speculated that other genetic mechanisms, possibly these other orthologous clusters, may function to generate H-MORPH in these strains. Therefore, we sought to determine the abundance of HAC, HBAC and H_C_AC throughout the *A. fumigatus* population. To do this we looked for the presence of *bafA* (HAC), *bafB* (HBAC), and *bafC* (H_C_AC) across available sequenced *A. fumigatus* strains. We confirmed that similar to *hrmA*, the number of strains positive for the presence of *bafA* is low (n=24) (Fig. 2). Similarly, *bafB* is present within ~27% of the *A. fumigatus* genomes analyzed (n=24) (Fig. 2). Strains positive for encoding *bafC* are more abundant (n=35), but this is complicated by the presence of another *bafC* ortholog in AF293 (Afu1g00770) that is not encoded near a putative *hrmC* ortholog (Fig. 2). Afu1g00700 shares 91% nucleotide identity with AFUB_096610 in CEA10 and while not syntenic, its high sequence similarity contributes to the positive identity of *bafC* in AF293 and potentially other strains as well (Fig. 2) (FungiDB) (33). Interestingly, there are strains similar to CEA10 that are positive for the presence of two or more of the *baf* genes (*bafA, bafB, bafC)* (n=21). However, this is likely an overestimate due to the presence of Afu1g00700 orthologs in some genomes.

**Figure 2.**
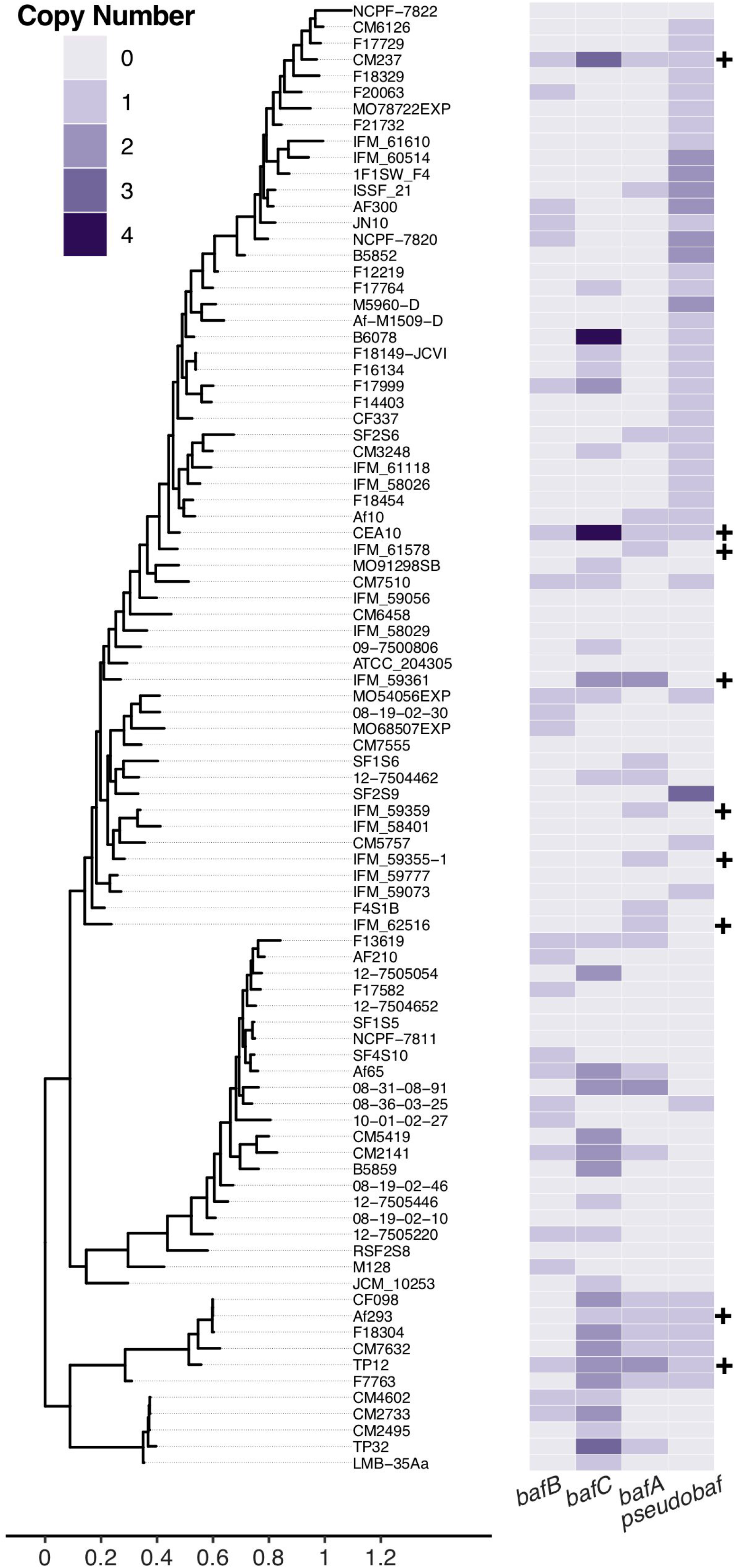
Phylogeny of 92 *A. fumigatus* strains with copy number of the cryptic gene and its putative orthologs. The *A. fumigatus* strain Maximum Likelihood phylogeny was constructed from 71,513 parsimony informative SNPs identified across the strains. The heat map indicates the abundance of *bafA, bafB, bafC*, and a *baf* pseudogene (*Pseudobaf*) across the phylogeny based on the genome sequences from CEA10. Strains which have been previously identified as encoding *hrmA* are indicated with a plus (+) sign.

Many of the analyzed genomes also encode a predicted pseudogene with high similarity to *bafB* (Pseudobaf) (Fig. 2). The pseudogenes are degenerate ORFs that have multiple stop codons throughout their sequence. Although AF293 does not encode *bafB* and *bafC* and their putative gene clusters, the presence of the pseudogene suggests that an ancestral strain of AF293 did encode *bafB* and HBAC. A BLAST search with *hrmB* or *bafB* from CEA10 against the AF293 genome matches a region of approximately 1,050 kb on Chromosome 3 where no genes are annotated (between Afu3g03760 and Afu3g03770) (Fig. S4A) (FungiDB) (33). The regions that map to *hrmB* and *bafB* are littered with stop codons truncating the ORFs, and thus are likely pseudogenes (Fig. S4B, C). If expressed, the chromosomal region that maps to *hrmB* in Af293 is predicted to generate a 123 amino acid protein instead of 423 amino acids (Fig. S4B); and the Pseudobaf chromosomal region that maps to *bafB* in AF293 is predicted to generate a 25 amino acid protein instead of 193 amino acids (Fig. S4C). The presence of other degraded ORFs, or pseudogenes, similar to *bafB* are observed across the phylogeny in different copy numbers (Fig. 2), posing interesting questions about potential functions of these pseudogenes, how they arose in the population, and how they are maintained.

### Introduction of the cryptic gene ortholog bafB is sufficient to complement the loss of cgnA and hrmA in EVOL20

To determine if *bafB* from CEA10, whose protein sequence is 78.35% identical to *bafA,* could complement the loss of *cgnA* in EVOL20 *(ΔcgnA^EVOL^),* we introduced *bafB* with the constitutively active *gpdA* promoter *(ΔcgnA^EVOL^; bafB^OE^).* The resulting strain reverted the N-MORPH phenotype of Δ*cgnA*^EVOL^ to the H-MORPH phenotype of EVOL20 with significantly increased colony furrows and percent vegetative mycelia (Fig. 3A, B). As mentioned above, the majority of HAC genes are not altered in expression as a result of *cgnA* deletion (26), thus the expression of other HAC genes could still be required for *bafB* to generate H-MORPH. The loss of *hrmA* in EVOL20 (Δ*hrmA*^EVOL^) reverts the colony to N-MORPH and mRNA levels of HAC genes are significantly reduced (26). To determine if *hrmA* and subsequently the HAC cluster genes that rely on *hrmA* for expression are necessary to generate H-MORPH in the presence of *bafB,* we introduced *bafB* with the constitutive *gpdA* promoter into the Δ*hrmA^EVOL^* strain (Δ*hrmA^EVOL^; bafB^OE^)*. Even in the absence of *hrmA, bafB* is sufficient to generate H-MORPH and significantly increase colony furrows and percent vegetative mycelia (Fig. 3A, B). In addition to H-MORPH, EVOL20 has elevated hypoxic fitness (H/N) and reduced surface adherence relative to AF293 that is dependent on both *hrmA* and *cgnA/bafA* (Fig. 1C, D) (26, 45). The overexpression of *bafB* significantly increases hypoxic fitness of Δ*hrmA^EVOL^* and Δ*cgnA^EVOL^* (Fig. 3C); and significantly reduces adherence of these strains to a plastic surface (Fig. 3D). Importantly, *bafB* is sufficient to complement these phenotypes in EVOL20 without increasing HAC gene mRNA levels (Fig. 3E). In fact, the mRNA levels of *hrmA* are slightly, but significantly, reduced as a result of constitutive *bafB* expression (Fig. 3E).

**Figure 3.**
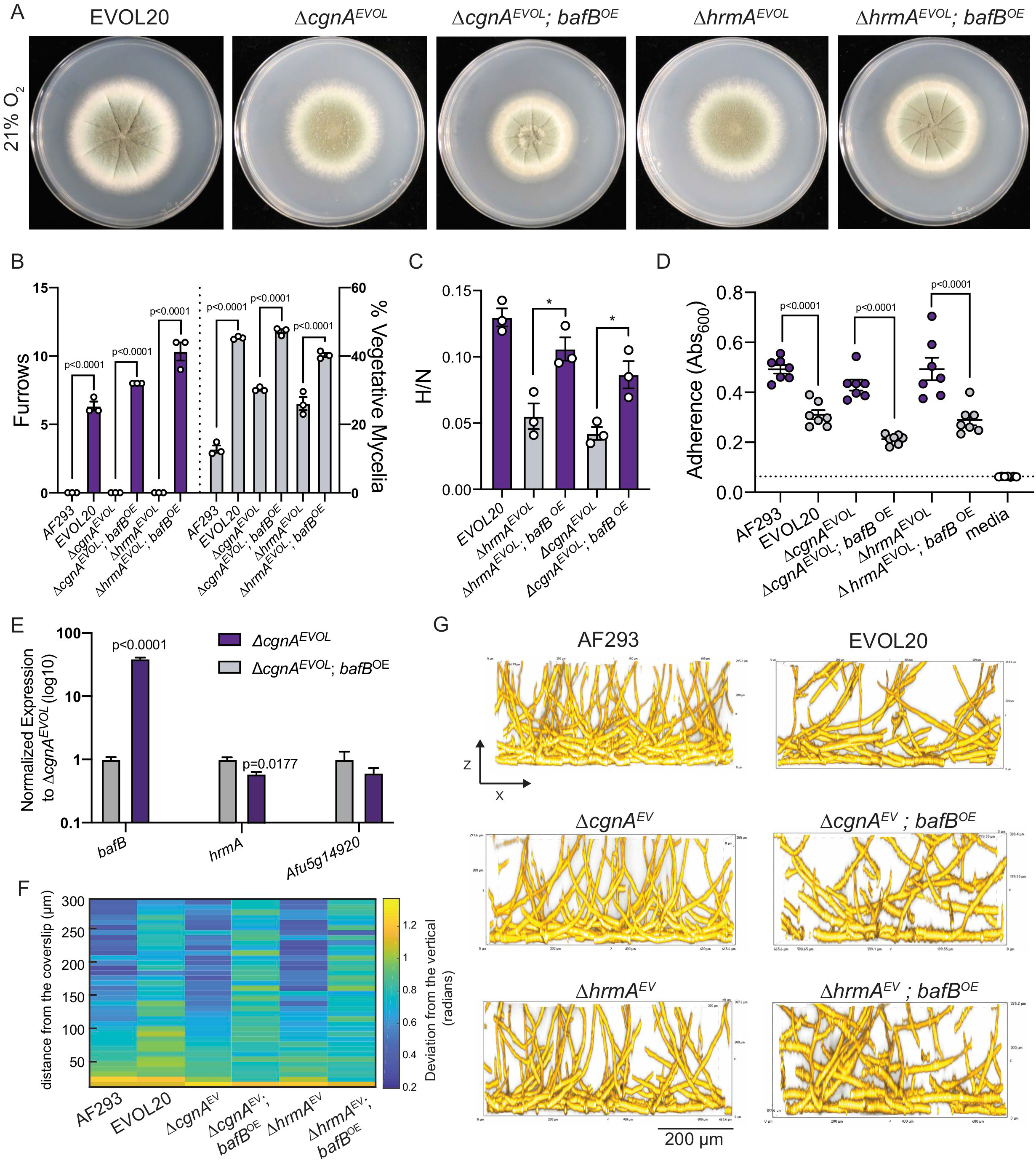
The putative ortholog of the cryptic gene, *bafB,* is sufficient to complement the loss of the HAC genes *cgnA* and *bafA.* (A) 96-hour colony biofilms from 21% O_2_ where hypoxia-locked (H-MORPH) morphological features, furrows and vegetative mycelia, can be visualized. Images are representative of three independent biological samples. (B) Quantification of the H-MORPH features from colony biofilms of three independent biological samples. Student’s two-tailed non-parametric t tests were performed between each isogenic strain set. (C) The ratio of fungal biomass in hypoxia (0.2% O_2_) relative to fungal biomass in normoxia (21% O_2_) (H/N) in shaking flask cultures. Student’s two-tailed non-parametric t test performed between isogenic strain sets. n=3 independent biological samples. (D) Adherence to plastic measured through a crystal violet assay. Dashed line marks the mean value for media alone. Student’s twotailed non-parametric t test performed between isogenic strain sets. n=7 independent biological samples. (E) Gene expression measured by qRT-PCR for representative HAC genes as a result of *bafB* overexpression at 21% O_2_. n=3 independent biological samples. (F) Heat map displaying the architecture of the fungal biofilms measured as the deviation of the hyphae from a vertical axis. Each column is representative of a minimum of three independent biological samples. (G) Representative images (n=3 biological samples) of submerged biofilms on the orthogonal plane (XZ) that are quantified in the heat map in (F). Scale bar is 200 μm. Error bars indicate standard error around the mean.

To test whether *bafB* expression alters biofilm architecture, a HAC-dependent phenotype of EVOL20, we cultured submerged biofilms for 24 hours and imaged the bottom ~300 μm of the biofilm. As a metric for biofilm architecture, we measured the angle of hyphal deviation from the vertical axis. As has been described for the N-MORPH strains AF293, Δ*cgnA^EVOL^,* and Δ*hrmA^EVOL^,* at 24 hours the bottom ~50 μm of the biofilm features filaments that grow along the surface and have a high deviation from the vertical (26). At depths above 50 μm for these N-MORPH strain, the hyphae orient vertically and grow polarized toward the air-liquid interface with little deviation from the vertical axis. In contrast, the H-MORPH strain EVOL20 features hyphae throughout all 300 μm that are oriented with a high deviation from the vertical (26). When *bafB* is overexpressed in the N-MORPH strains *cgnA^EVOL^* and Δ*hrmA^EVOL^,* the resulting H-MORPH strains (Fig. 3A) develop biofilms that also resemble the architecture of EVOL20 (Fig. 3 F, G). There is greater hyphal deviation from the vertical axis above 50 μm in the biofilms of Δ*cgnA^EVOL^; bafB^OE^* and Δ*hrmA^EVOL^; bafB^OE^* (Fig. 3F, G). Thus, introduction of a constitutively expressed *bafB* is sufficient to complement the HAC-dependent phenotypes of EVOL20.

The mechanisms underlying hyphal arrangement and ultimately the shift from vertically-oriented hyphal growth to a more horizontal hyphal growth in the biofilm remains undefined. Previously we had hypothesized that this was a consequence of the altered hyphal surface of H-MORPH strains (26). The BafB protein is predicted to have a signal sequence at its N-terminus (SignalP, FungiDB) (Fig. S5A) (33, 46). To gain insight into how *bafB* could directly impact the biofilm architecture of the Δ*cgnA^EVOL^* strain, we generated a c-terminal green fluorescent protein (GFP) tagged allele of *bafB* in Δ*cgnA^EVOL^.* Introduction of the GFP-tagged allele, like the native *bafB* allele, is able to revert the N-MORPH colony morphotype of Δ*cgnA^EVOL^* to H-MORPH (Fig. S5B). In mature hyphae, the localization of the GFP signal is present both in the cytosol within circular structures that resemble trafficking endosomes or vacuoles previously described in *A. nidulans* (47) (Fig. S5C), and concentrated toward the distal hyphal region (Fig. S5D). At the distal region, the GFP signal is present within circular structures, or puncta, as well as localized along the sides of the hyphae (Fig. S5D). Time-lapse imaging reveals that these BafB puncta are dynamic and move rapidly within the hyphae (Video S1, Fig. S5E). Co-staining with the membrane dye FM4-64 indicate overlap in the patterns of BafB localization and endosome localization (Fig. S5F). This subcellular pattern and the presence of the N-terminal secretion signal peptide (Fig. S5A) support to the hypothesis that BafB localizes extracellularly at the hyphal tips or is secreted (48). While the GFP signal corresponding to BafB is largely absent from the hyphal edges where the cell wall is more stable, the signal is abundant at the tip where cell wall modeling is in progress. This is evidenced by the absence of Dectin-1 binding, which specifically interacts with β-1,3-glucan, at the hyphal tip where BafB is abundant (Fig. S5G). Cell wall irregularities are a feature of H-MORPH and the actively growing hyphal tip directs cell polarity (26, 49). As colony morphology is a consequence of polarized growth and structure of the cell wall this localization pattern indicates that BafB could be acting as the H-MORPH effector (26, 50). The high amino acid identity shared between *bafB* and the HAC-resident gene *bafA* raise the question of whether *bafA* is the HAC effector and is sufficient to generate H-MORPH in the parental strain AF293.

### Overexpression of bafA generates H-MORPH and elevated hypoxic growth in the absence of HAC induction in two independent strain backgrounds

In the parental strain AF293, the basal expression of HAC is low, and previous RNA-sequencing data reveals no mapped reads to the predicted *bafA* ORF in AF293 (Fig. S2A) (26). In addition, qRT-PCR for *bafA* mRNA revealed no detection above background in AF293, but overexpression of an additional *bafA* allele results in detectable *bafA* mRNA (Fig. S6A). The synthetic, elevated expression of *bafA* in AF293 results in H-MORPH colony morphology with significantly increased colony furrows and percent vegetative mycelia relative to AF293 (Fig. 4A, C). Interestingly, the colony morphology in hypoxia (0.2%O_2_) is also distinctly different as a result of *bafA* overexpression. Unlike AF293, the colony in hypoxia is small, dense and lacks furrows and conidiation (Fig. 4A), resembling the previously published colony morphology resulting from constitutive *hrmA* expression (26).

**Figure 4.**
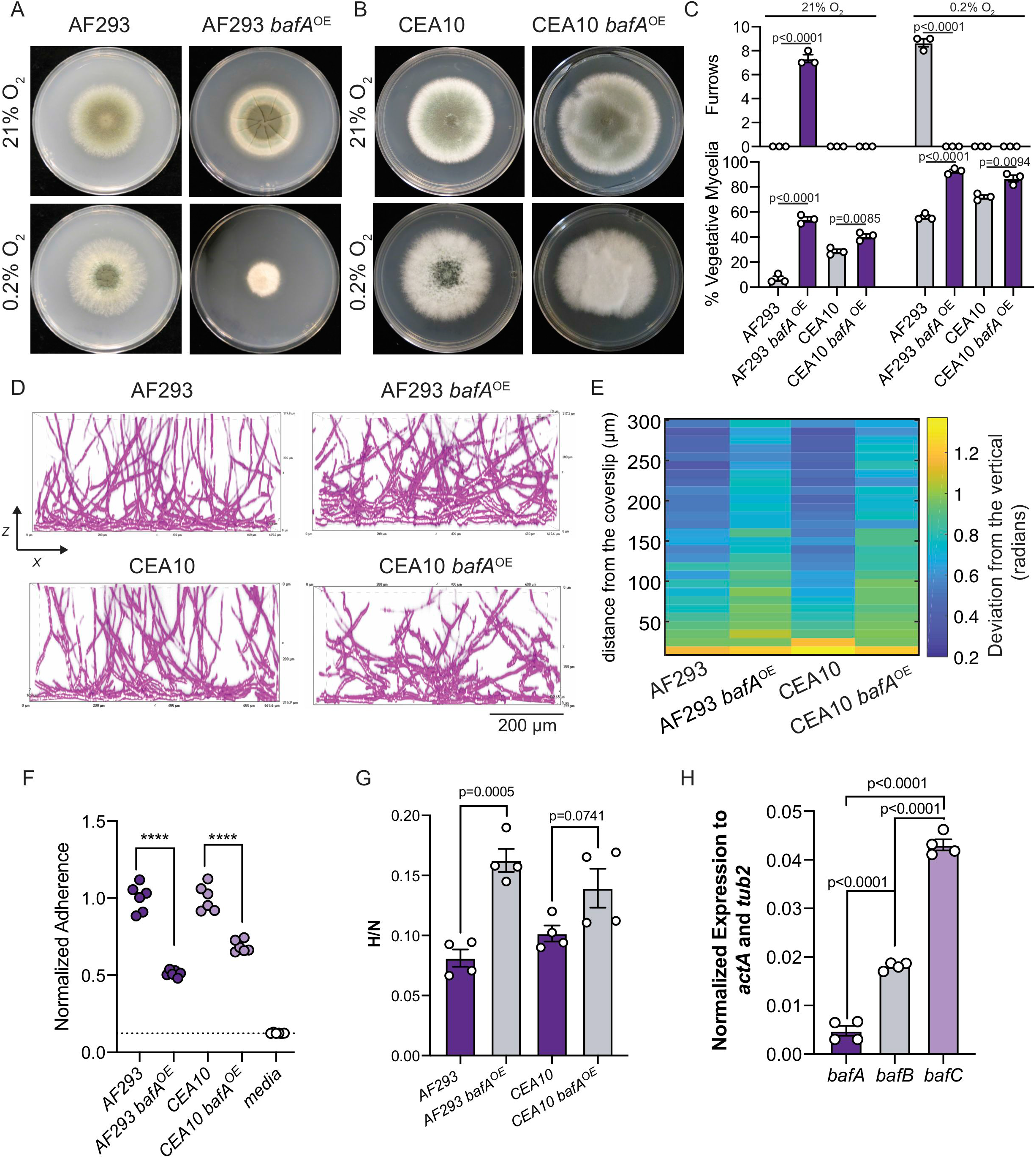
Introduction of the HAC cryptic gene *bafA* is sufficient to generate H-MORPH in AF293 and impacts biofilm architecture in the baf^+^ strain CEA10. (A) 96-hour colony biofilms in normoxia (21% O_2_) and hypoxia (0.2% O_2_) of AF293 and AF293 with the overexpression of *bafA*. Images are representative of three independent biological samples. (B) 96-hour colony biofilms in normoxia (21% O_2_) and hypoxia (0.2% O_2_) of CEA10 and CEA10 with the overexpression of *bafA.* Images are representative of three independent biological samples. (C) Quantification of the H-MORPH features from colony biofilms of three independent biological samples. Student’s two-tailed nonparametric t tests were performed between each isogenic strain set. (D) Representative images of submerged biofilms (n=3 biological samples) on the orthogonal plane (XZ). Scale bar is 200 μm. (E) Heat map displaying the architecture of the fungal biofilms measured as the deviation of the hyphae from a vertical axis. Each column is representative of a minimum of three independent biological samples. (F) Adherence to plastic measured through a crystal violet assay. Dashed line marks the mean value for media alone. Student’s two-tailed non-parametric t test performed between isogenic strain sets. n=6 independent biological samples. (G) The ratio of fungal biomass in hypoxia (0.2% O_2_) relative to fungal biomass in normoxia (21% O_2_) (H/N) in shaking flask cultures. Student’s two-tailed non-parametric t test performed between isogenic strain sets. n=4 independent biological samples. (H) Gene expression measured by qRT-PCR for *bafA*, *bafB,* and *bafC* in AF293 and CEA10. n=4 independent biological samples. Error bars indicate standard error around the mean.

The strain CEA10 contains HAC, HBAC, and H_C_AC, but like AF293, *bafA* expression is below the level of detection by qRT-PCR in biofilm cultures but can be detected following introduction of a second overexpressed *bafA* allele (Fig. S6B). Elevated expression of *bafA* in CEA10 qualitatively alters the colony morphology in normal (21% O_2_) and low oxygen (0.2% O_2_) and significantly increases the percent vegetative mycelia (Fig. 4B, C). However, no colony furrows are present as a result of *bafA* constitutive expression in CEA10 (Fig. 4C). Despite the absence of this macroscopic H-MORPH feature, overexpression of *bafA* in CEA10, and in AF293, impacts biofilm architecture by increasing the deviation of hyphae from the vertical axis above the bottom 50 μm of the biofilm (Fig. 4D, E). Unlike AF293, even during hypoxic growth CEA10 colonies do not feature furrows, and instead abundant aerial hyphae develop generating a ‘fluffy’ colony morphotype. We speculate that perhaps there is a dichotomy among strains of *A. fumigatus* where some respond to low oxygen by forming aerial hyphae (i.e. CEA10) and others develop furrows (i.e. AF293).

H-MORPH in EVOL20, and other clinical isolates, coincides with reduced adherence and increased hypoxic fitness (hypoxic growth relative to normoxia growth, H/N) (26). In both CEA10 and AF293, overexpression of *bafA* significantly reduces hyphal adherence to plastic (Fig. 4F). Despite documented differences in hypoxic growth between AF293 and CEA10, *bafA* overexpression also significantly increases the hypoxic fitness of both strains, though to a lesser extent in CEA10 (Fig. 4G) (45). The inability for *bafA* expression to impact CEA10 colony morphology, and its apparent reduced impact on adherence and hypoxic growth relative to AF293 may be explained by the presence of the other *baf* genes encoded in the CEA10 genome. While *bafA* mRNA levels are undetectable in CEA10 during normal oxygen growth, mRNA for both *bafB* and *bafC* is detected (Fig. 4H). As the amino acid identity between these three proteins ranges from 45-78% we hypothesize that *bafB* and *bafC* are also sufficient to impact colony and biofilm morphology.

### Overexpression of the bafA orthologs bafB and bafC generate H-MORPH-like phenotypes and impact hypoxic growth

To determine if *bafB* and *bafC* are sufficient to generate H-MORPH phenotypes in the independent reference strains AF293 and CEA10, we used a constitutive promoter to drive expression of these genes and assessed colony morphology, adherence, and biofilm architecture. Introduction of either *bafB* or *bafC* in AF293 generates features of H-MORPH in normoxia with significantly increased furrows and percent vegetative mycelia (Fig. 5A, C). Similar to *bafA* overexpression in CEA10, *bafB* overexpression did not induce H-MORPH features of colony furrows and increased percent vegetative mycelia in CEA10 (Fig. 5B, D). However, *bafB* expression significantly reduced overall conidiation in normoxia (21% O_2_) and hypoxia (0.2% O_2_), a complimentary metric to percent vegetative mycelia (Fig. 5E). Overexpression of *bafC* in CEA10 is unique in that it does significantly increase colony furrows in normoxia relative to CEA10 (Fig. 5B, D). However, the percent vegetative mycelia is not significantly increased (Fig. 5D).

**Figure 5.**
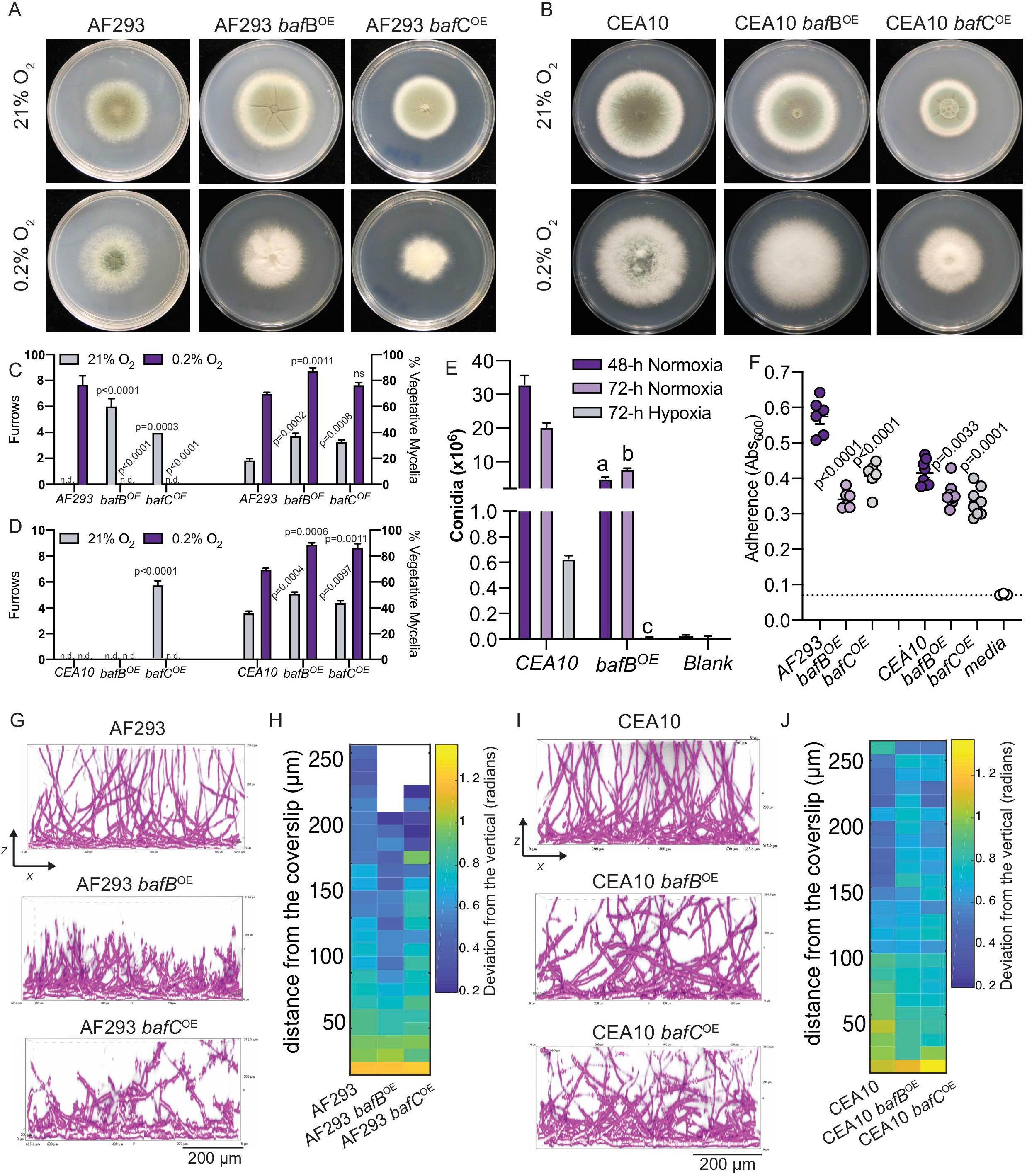
Introduction of *bafB* and *bafC* impact colony and submerged biofilm morphology in independent strain backgrounds. **(A)** 96-hour colony biofilms in normoxia (21% O_2_) and hypoxia (0.2% O_2_) of AF293 and AF293 with the overexpression of *bafB* or *bafC.* Images are representative of three independent biological samples. (B) 96-hour colony biofilms in normoxia (21% O_2_) and hypoxia (0.2% O_2_) of CEA10 and CEA10 with the overexpression of *bafB* or *bafC.* Images are representative of three independent biological samples. (C) Quantification of the H-MORPH features from colony biofilms of AF293, AF293 *bafB^OE^*, and AF293 *bafC^OE^* with three independent biological samples. One-way ANOVA with Dunnett’s posttest for multiple comparisons relative to AF293 were performed. (D) Quantification of the H-MORPH features from colony biofilms of CEA10, CEA10 *bafB^OE^,* and CEA10 *bafC^OE^* with three independent biological samples. One-way ANOVA with Dunnett’s posttest for multiple comparisons relative to CEA10 were performed. (E) Quantification of conidiation from three independent biological samples of CEA10 and CEA10 *bafB^OE^* in normoxia (21% O_2_) or hypoxia (0.2% O_2_). Student’s two-tailed non parametric t tests were performed between CEA10 and CEA10 *bafB^OE^* for each time point. (a: p=0.0004, b: p=0.0006, c: p<0.0001). (F) Adherence to plastic measured through a crystal violet assay. Dashed line marks the mean value for media alone. One-way ANOVA with Dunnett’s posttest for multiple comparisons was performed between isogenic strain sets relative to AF293 or CEA10. n=6 independent biological samples for AF293 strains and n=8 independent biological samples for CEA10 strains. (G) Representative images of submerged biofilms (n=3 biological samples) on the orthogonal plane (XZ) of AF293, AF293 *bafB^OE^*, and AF293 *bafC^OE^*. Scale bar is 200 μm. (G) Representative images of submerged biofilms (n=3 biological samples) on the orthogonal plane (XZ) of CEA10, CEA10 *bafB^OE^*, and CEA10 *bafC^OE^.* Scale bar is 200 μm. (I) Heat map displaying the architecture of the fungal biofilms measured as the deviation of the hyphae from a vertical axis. Each column is representative of a minimum of three independent biological samples.

Despite variation in how the *baf* genes impact colony morphology in the two strain backgrounds, in both AF293 and CEA10 overexpression of *bafB* or *bafC* results in significantly reduced adherence to plastic (Fig. 5F). CEA10 adheres less well to plastic compared to AF293, and the difference in adherence is smaller as a result of *bafB* or *bafC* overexpression. As these two genes are already present and expressed in CEA10 (Fig. 4H), it is possible that this native *baf* expression contributes to this difference between CEA10 and AF293.

As putative biofilm architecture factors, we sought to confirm an impact of *bafB* and *bafC* on biofilm architecture, similar to that observed with elevated expression of *bafA* (Fig. 3D, E). In AF293, overexpression of *bafB* visibly impacts biofilm architecture and formation in the XZ (Fig. 5G) and XY dimensions (Fig. S6C). The XY dimension reveals dense hyphal growth and abundant hyphal branching (Fig. S6C). The XZ dimension shows a stunted 24 hour biofilm that reaches heights of only 200-250 μm (Fig. 5G). Similarly, regions of the 24 hour biofilms generated by the overexpression of *bafC* in AF293 (AF293 *bafC^OE^*) are also stunted with evidence of hyphae that are hyper branching (Fig. 5G, Fig. S6C). In regards to biofilm architecture as defined by hyphal orientation to the vertical axis, overexpression of *bafC* but not *bafB* in AF293 results in increased deviation from the vertical axis above 50 μm (Fig. 5 G. H). Notably, constitutive expression of *bafA*, *bafB*, or *bafC* in AF293 also impacts morphology during liquid growth similar to that of EVOL20 (Fig. S6D).

In CEA10 biofilms, overexpression of *bafB* and *bafC* results in increased deviation from the vertical axis above 50 μm in 24 hour biofilms (Fig. 5I, J). There is also qualitative evidence for hyper branching as a result of elevated *bafB* or *bafC* expression in CEA10 (Fig. S6E). These data support a role for all three proposed *baf* genes in biofilm architecture, through multiple metrics, in two independent strain backgrounds of *A. fumigatus.*

### Introduction of A. fumigatus bafA into Aspergillus niger generates H-MORPH and simultaneously increases hypoxic growth

We have previously reported that among the *Aspergilli*, *hrmA* is absent from the notable species of *A. nidulans, A. oryzae, A. flavus,* and *A. niger* based on available genome sequences (26). However, *Aspergillus niger* strain CBS 513.88 encodes a gene, An08g12010, with 69% nucleotide identity to *A. fumigatus bafA* and 41.03% amino acid identity to the predicted protein sequence of BafA (Fig. S7A). This suggests that the role of *baf* or *baf-like* genes may be conserved in other *Aspergillus* species. We sought to determine if *A. fumigatus bafA (AfbafA)* could influence colony morphology, biofilm architecture, hypoxic growth, and adherence in the *A. niger* reference strain A1144. This strain was selected for its robust growth at 37° and the ease at which it is genetically manipulated.

We overexpressed *A. fumigatus bafA* in *A. niger* with the constitutive *gpdA* promoter to generate An *AfbafA^OE^* (Fig. S7B). Overexpression of *bafA* in *A. niger* generated H-MORPH colonies with significantly increased colony furrows and percent vegetative mycelia compared to the control A1144 (Fig. 6A, B). Intriguingly, the overexpression of *AfbafA* in *A. niger* resulted in the production of a bright yellow pigment, shown here in two independent transformants (Fig. 6A). The production of yellow pigments by *A. niger* has been noted in the literature for decades as a result of various growth conditions and genetic manipulations (51).

**Figure 6.**
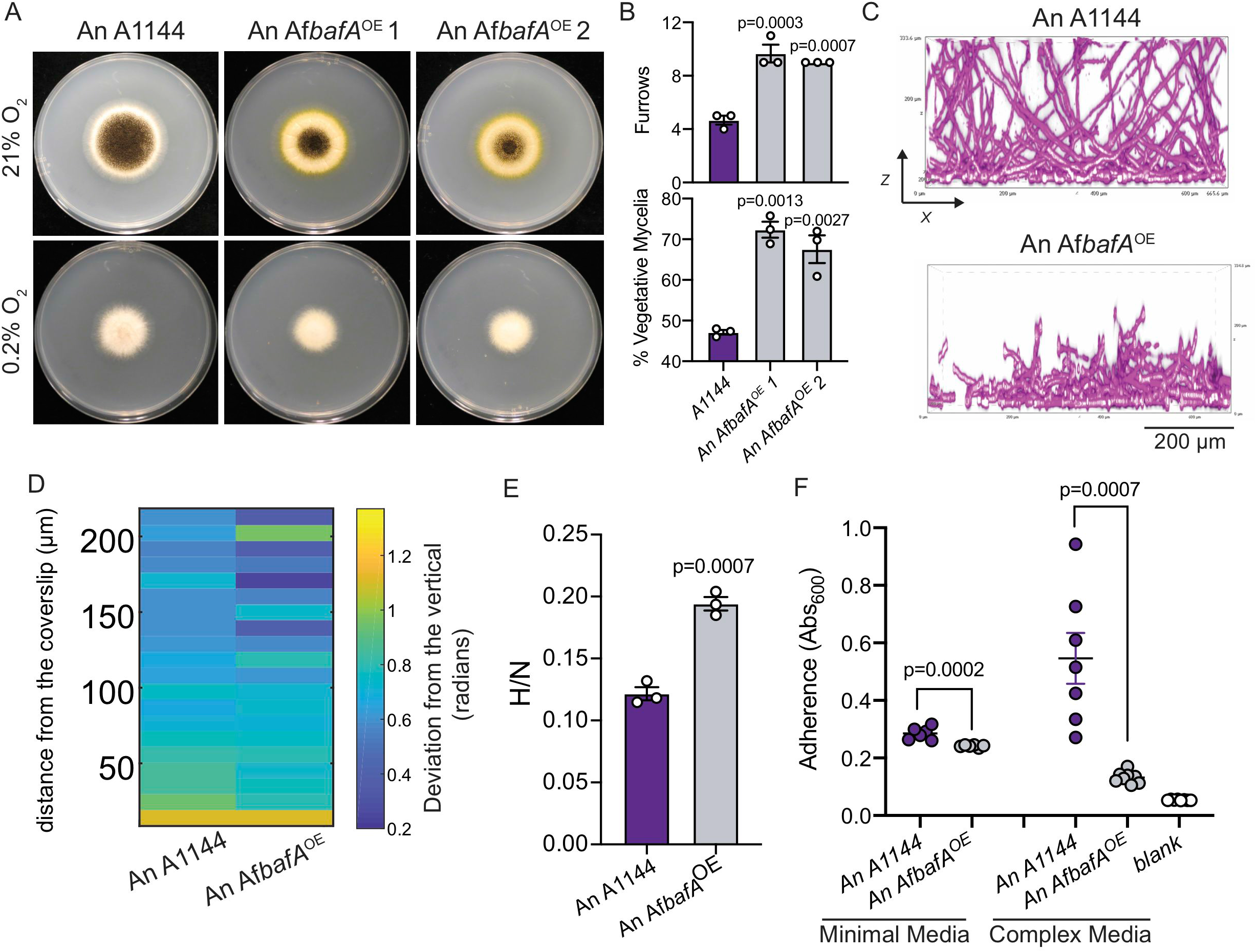
Introduction of *A. fumigatus bafA* is sufficient to generate H-MORPH in *Aspergillus niger*. (A) 96-hour colony biofilms in normoxia (21% O_2_) and hypoxia (0.2% O_2_) of *A. niger* reference strain A1144 and two independent strains of A1144 with the overexpression of *A. fumigatus bafA (AfbafA^OE^).* Images are representative of three independent biological samples. (B) Quantification of the H-MORPH features from colony biofilms of three independent biological samples in normoxia (21% O_2_). One-way ANOVA with Dunnett’s posttest for multiple comparisons relative to A1144. (C) Representative images of submerged biofilms (n=3 biological samples) on the orthogonal plane (XZ). Scale bar is 200 μm. (D) Heat map displaying the architecture of the fungal biofilms measured as the deviation of the hyphae from a vertical axis. Each column is representative of a minimum of three independent biological samples. (E) The ratio of fungal biomass in hypoxia (0.2% O_2_) relative to fungal biomass in normoxia (21% O_2_) (H/N) in shaking flask cultures. Student’s two-tailed non-parametric t test performed between isogenic strain sets. n=3 independent biological samples. (F) Adherence to plastic measured through a crystal violet assay. Student’s two-tailed non-parametric t test performed between samples within each media type. n=6 independent biological samples for minimal media and n=7 independent biological samples for complex media (minimal media with yeast extract).

Similar to *A. fumigatus,* the *A. niger* reference strain A1144 forms a submerged biofilm with dense filaments within the first 50 μm that are oriented perpendicular to the vertical axis (Fig. 6C, D). Above the ~50 μm at the base of the biofilm, filaments become oriented more closely along the vertical axis, similar to what has been observed with N-MORPH strains of *A. fumigatus* (i.e. AF293) (Fig. 6C, D). Introduction of the constitutively expressed *AfbafA* alters the biofilm of A1144. At 24 hours, the hyphae are stunted reaching heights of only 200-250 μm in height (Fig. 6C). These stunted filaments highly deviate from the vertical axis throughout the height of the biofilm indicating that *AfbafA* is capable of impacting biofilm architecture across fungal species (Fig. 6D).

Not only does *AfbafA* impact the colony morphology to generate H-MORPH and modulate the biofilm architecture, but it also generates other H-MORPH and EVOL20 associated phenotypes including increased hypoxia fitness and reduced adherence. In AF293 and CEA10 expression of *bafA* results in increased hypoxia fitness (hypoxic growth normalized to normoxic growth); similarly, the hypoxia fitness of A1144 significantly increases with constitutive expression of *AfbafA* (Fig. 6E). Adherence of *A. fumigatus* is quantified in minimal media, however, the adherence of the reference *A. niger* strain A1144 is low in minimal media. Thus, we opted to quantify the impact of *AfbafA* on *A. niger* adherence in both minimal and complex media where A1144 adherence is more robust. In both conditions, adherence was significantly reduced with expression of *AfbafA* compared to A1144 (Fig. 6F). Not only do reduced adherence and increased hypoxia fitness track with H-MORPH on the macroscale and microscale, as has been observed previously, but they do so as a result of *bafA* expression across different *Aspergillus* species. The ability of *bafA* alone to generate these phenotypes in the two independent species of *Aspergillus* supports its role as the effector protein of HAC, and supports its application to modify biofilm architecture and function in *Aspergillus* species.

## Discussion

Herein we identify and characterize a family of putative orthologous proteincoding genes that are heterogeneously expressed across *A. fumigatus* strains and impact biofilm architecture and hypoxic growth. Biofilm architecture refers to the complex arrangement of cells within a three dimensional structure that develops after initial surface attachment and monolayer growth (52). The architecture of the microbial biofilm is dependent on the organism, the surface, and the exogenous environment (52). Specific biofilm architectures have been associated with tolerance to desiccation and antibiotics (53–55), phage resistance (56), and predation evasion (57). Examples of biofilm architecture for bacterial biofilms include the formation of pillars and mushroomlike structures (52, 58, 59). For fungi, biofilm architecture takes on additional dimensions of complexity. Biofilms of the polymorphic yeast *C. albicans* reflect the architectural arrangement of multiple cell morphologies (60); and for the filamentous fungi, directional growth of multicellular hyphae and hyphal branching complicate biofilm architecture (26). The important ecological, clinical, and industrial roles of biofilms, and the relationship between biofilm structure and function has stimulated the characterization of molecular components that influence biofilm architecture (61, 62). Such factors have most thoroughly been studied in bacteria and yeast (63–66). Filamentous fungi form mycelia, or biofilms, with intricate hyphal architectures, yet similar molecular components remain ill defined. We begin to address this gap in knowledge through characterization of the heterogeneous *baf* gene family in *A. fumigatus.*

All three genes, *bafA*, *bafB*, and *bafC* are previously uncharacterized and *bafB* and *bafC* are annotated as encoding hypothetical proteins. With no conserved protein domains or characterized homologs, it is difficult to ascertain the putative molecular functions of these proteins. Preliminary phylogenetic analyses suggest these genes are restricted to the Eurotiales and rare outside the *Aspergillus* genus. Microscopy studies with a GFP-tagged BafB, which shares 78.35% amino acid identity with the predicted BafA protein sequence, reveals a localization pattern suggesting BafB is transported to the distal region of the hyphae within endosomes, and concentrates at the growing tip (Fig. S5). The BafB protein does have a predicted N-terminus secretion signal and further experiments are underway to investigate if BafB is secreted from the hyphae or localized extracellularly. The observed sub-cellular localization of BafB is intriguing because the distal region of the hyphae is where polarized growth is regulated, cell wall synthesis takes place, and protein secretion occurs (48, 67). In a HAC induced H-MORPH strain, the hyphae exhibit a modified cell wall and a biofilm architecture phenotype, where hyphae grow horizontally and no longer polarize to the same degrees along the vertical axis (26). One appealing hypothesis is that BafB acts at the hyphal tip and surrounding distal region where the rigid cell wall is yet to be constructed to directly modify these pathways (Fig. 6SG). Alternatively, we cannot rule out that these phenotypes are the result of BafB modulating the secretory pathway, resulting in diverse downstream phenotypes dependent on secretion (polarized growth, cell wall synthesis, etc.). Disruption of *A. fumigatus* membrane trafficking through the deletion of the Rab GTPase *sec4* homolog *srgA,* does result in unstable and diverse colony morphologies (68). Work is ongoing to determine the molecular function of these important proteins and how they impact H-MORPH and related phenotypes.

To-date, surface adherence of *A. fumigatus* is attributed to the production of the primary exopolysaccharide galactosaminogalactan (GAG), a polysaccharide also produced by *A. niger* (69, 70). Microscopy of EVOL20 H-MORPH biofilms reveals a reduction in hyphal-attached polysaccharide compared to AF293 which is reflected in the reduced adherence of EVOL20 to plastic relative to AF293 (26). The matrix detachment is credited to the altered cell surface of H-MORPH strains. As introduction and constitutive expression of *bafA* in *A. fumigatus* and *A. niger* reduces the surface adherence characteristic of H-MORPH, it is likely modifying the hyphal surface similar to EVOL20. This modification of the surface could be due to BafA localizing to the surface and directly preventing GAG attachment, or BafA could modify the hyphal surface indirectly through regulation of cell wall synthesis and protein/carbohydrate secretion. We do not know how GAG attaches to the *A. fumigatus* hyphal surface and if adherence is mediated by polysaccharides or proteins, but both models would be consistent with the localization patterns we observe with the highly similar BafB at the distal hyphal region.

H-MORPH, as the name implies, is also tightly associated with *A. fumigatus* hypoxic growth. The colony morphology features characteristic of H-MORPH, colony furrows and vegetative, non-conidiating mycelia, are hallmarks of many, but not all, *A. fumigatus* colonies grown in low oxygen environments (26). The EVOL20 strain, which was serially passaged in low oxygen, has significantly increased hypoxic growth compared to the parental, pre-passaged strain AF293 (26, 45). This increased hypoxic growth of EVOL20 is dependent on *bafA*, and *bafA* is sufficient to increase the hypoxic growth of *A. fumigatus* and *A. niger* when it is constitutively expressed. How H-MORPH hyphae facilitate increased growth in low oxygen remains unknown, and is the focus of ongoing work. The formation of wrinkles in bacteria and yeast colony biofilms have been associated with increased oxygen penetration in the biofilm (24, 71) and an altered redox state (72, 73). We hypothesize that the H-MORPH hyphal surface allows for the formation of furrows that potentially increase the colony surface area exposed to ambient oxygen.

The formation of aerial hyphae and the generation of a ‘fluffy’ colony morphology would also increase the hyphal surface area exposed to ambient oxygen. In surface colonies of *A. oryzae*, oxygen levels remain high within the entire 4 mm of aerial hyphal growth and drop precipitously at the dense mycelial base (74). The generation of aerial hyphae is a phenotype associated with hypoxic colony growth of the reference strains CEA10 and AfS35 (31, 75). However, this is not the case with the reference strain AF293, where instead the colonies form furrows in response to low oxygen (26). As noted in our data above, CEA10 does not form furrows during low oxygen growth (Fig. 4B). Our lab and others have described phenotypic differences between AF293 and CEA10 as examples of the natural heterogeneity within the *A. fumigatus* species (45, 76–78). An interpretation is there is a phenotypic dichotomy within *A. fumigatus* separating those strains that form aerial hyphae and ‘fluffy’ colonies in low oxygen and those that form furrows in low oxygen. The heterogeneity of *baf* gene presence or absence across the strain phylogeny could facilitate the strain-specific morphological adaptation to hypoxia. In support of this hypothesis, constitutive expression of *bafC* in CEA10 is able to generate colony furrows in normal oxygen (21% O_2_) and simultaneously impact the pattern of aerial hyphae production (‘fluffiness’) during low oxygen growth (Fig. 5B). Currently a lack of morphological data, particularly in low oxygen, for publicly available genome-sequenced strains of *A. fumigatus* limits our ability to comprehensively interrogate the relationship between low oxygen colony growth strategies and *baf* gene function, but work to address this is ongoing. Simultaneously, future work is focused on quantifying aerial hyphae production in *A. fumigatus* colonies, a morphology we suspect represents a second variant of H-MORPH.

Surface adherence and low oxygen growth are both intimately related to *A. fumigatus* pathogenesis. The polysaccharide galactosaminogalactan has known immune-modulatory effects (79) and its loss increases exposure of inflammatory ß-glucans (69). An *A. fumigatus s*train that does not produce galactosaminogalactan is attenuated in virulence in multiple murine models of invasive aspergillosis (69). *A. fumigatus* requires the ability to adapt to low oxygen to cause disease (31) and low oxygen growth *in vitro* correlates with virulence (45). The ability to synthetically modulate or target these cellular processes in addition to altering biofilm architecture *in vivo* could have important implications in the treatment of disease caused by *A. fumigatus*. Targeted manipulation of lesion architecture could increase drug permeability or increase oxygen permeation within biofilms to alter the fungal physiology and host response. The *baf* proteins influence on biofilm architecture, adherence, and hypoxic growth warrant further investigation to evaluate their potential roles as adjunctive therapeutic targets. Additionally, biofilms are a staple of *Aspergillus* industrial processes, where their presence can be beneficial or detrimental to product yield. Surface-immobilized biofilms can greatly increase product yields but can also be damaging and corrosive to industrial materials (19, 20). Targeted genetic manipulation of industrial strains for the purposes of enhancing or reducing biofilm growth, modifying biofilm architecture, or triggering biofilm detachment requires known genetic components, of which the *baf* genes are prime potential candidates.

## Acknowledgments

We thank A. Lavanway (Dartmouth) for her microscopy expertise. We thank Dr. Isabelle Benoit Gelber (Concordia Canada) for advice on genetic manipulation of *A. niger.* This work was supported by the efforts of R.A.C through funding by NIH National Institute of Allergy and Infectious Diseases (NIAID) (grant nos. R01AI130128 and R01AI146121). R.A.C holds an Investigators in Pathogenesis of Infectious Diseases Award from the Burroughs Wellcome Fund. C.H.K. was supported by the NIH NIAID Ruth L. Kirschstein National Research Service Award (no. F31AI138354). JES is CIFAR Fellow in the program Fungal Kingdom: Threats and Opportunities. CDN is supported by the National Science Foundation grant MCB 1817342, a Burke Award from Dartmouth College, a pilot award from the Cystic Fibrosis Foundation (STANTO15RO), NIH grant P30-DK117469, NIH grant P20-GM113132 to the Dartmouth BioMT COBRE, and grant RGY0077/2020 from the Human Frontier Science Foundation with co-PI Alexandre Persat.

## Supplemental Material Legends

**Figure S1. H-MORPH associated phenotypes and the HAC gene cluster.** (A) N-MORPH (AF293) grown at 21% O_2_ and H-MORPH (EVOL20) grown at 21% O_2_. Black arrow indicates furrows and purple arrow indicates the perimeter of vegetative mycelia (PVM) as defined previously (26). (B) Within the subtelomeric region of chromosome 5 in the AF293 genome seven annotated genes belong to the *hrmA-associated* gene cluster (HAC). The putative regulator is *hrmA* and the annotated *cgnA* coding sequence were demonstrated to be essential for H-MORPH in EVOL20. (C) Strains previously described by Kowalski et al. (26) and their associated relevant phenotypes.

**Figure S2. Deletion of *cgnA* in two independent strain backgrounds reduces mRNA expression the cryptic ORF.** (A) Representative alignment of RNA-sequencing reads from Kowalski et al. 2019 (26) in IGV: Integrative Genomics Viewer within the HAC region. The green dashed box indicates mapped reads from the EVOL20 sample where no annotated gene is present. (B) Gene expression of *cgnA* and the cryptic ORF quantified through qRT-PCR in EVOL20 and Δ*cgnA*^EVOL^ in normoxia (21% O_2_). n=3 independent biological samples. Student’s two-tailed non-parametric t test was performed between each strain for each gene. (C) Gene expression of *cgnA* and the cryptic ORF quantified through qRT-PCR in AF293, *hrmA^R-EV^* and *hrmA*^R-EV^; Δ*cgnA* in normoxia (21% O_2_). n=3 independent biological samples. One-way ANOVA with Tukey’s multiple comparison test was performed for each gene. Error bars indicate standard error around the mean. (D) The predicted exons and introns that correspond to the cryptic ORF within HAC from NCBI ORF Finder. (E) Schematics of DNA constructs utilized in generating the *cgnA^OE^* and *cgnA^RECON^* strains in the Δ*cgnA^EVOL^* strain. The sequence for the *gpdA* promoter and *trpC* terminator are from *A. nidulans.*

**Figure S3. CEA10 orthologous sequences to the HAC cryptic gene.** (A) Schematic alignment of the hrmA associated gene cluster (HAC) and the putative orthologous gene clusters identified in the strain CEA10. Grey boxes align putative orthologous genes. Not drawn to scale. (B) The protein sequence alignment between the cryptic gene BafA (Predicted) and CEA10 BafB (AFUB_044360). The sequences share 78.35% identity. (C) The protein sequence alignment between the cryptic gene BafA (Predicted) and CEA10 BafC (AFUB_096610). The sequences share 45.41% identity.

**Figure S4. CEA10 HBAC maps to a region of degraded ORFs in AF293.** (A) The loci on AF293 chromosome 3 where the HBAC genes have high nucleotide (nt) identity. (B) Translation of the degraded ORF in AF293 that shares high similarity with *hrmB* from CEA10. The region corresponding to the *hrmB* intron was spliced out before translation. (C) Translation of the degraded ORF in AF293 that shares high similarity with *bafB* from CEA10. The region corresponding to the *bafB* intron was spliced out before translation. Yellow boxes and asterisks indicate stop codons within the translated degraded ORFs.

**Figure S5. A GFP-tagged allele of *bafB* reveals BafB localizes within endosomes at the distal hyphal region.** (A) A protein model for BafB with the SignalP 5.0 predicated signal peptide from amino acids (AA) 0 – 21 with an NN D-Score of 0.592 based on both SP-NN and SP-HMM algorithms with a 0.75 HMM signal probability. (B) A c-terminal GFP tagged allele of *bafB* generates H-MORPH in the N-MORPH strain Δ*cgnA^EVOL^.* (C) A representative hypha revealing GFP signal present within the cytosol in round cellular structures and concentrated at the distal end of the filament associated with the inner or outer surface. Inset panel shows magnification of the hyphal tip. Images are representative of a minimum of ten independent biological samples. Scale bar is 20 μm. (D) Magnified images of two hyphal tips revealing the concentration of GFP signal at the tip in greater detail. Images are presentative of a minimum of ten independent biological replicates. Scale bar is 10 μm. (E) Still frames from Video S1 of Δ*cgnA*^EVOL^; *bafB*^OE-GFP^ where GFP signal is shown in black. Puncta of BafB can be seen moving between frames (purple and black arrows). (F) FM4-64 localized in similar patterns as BafB at the hyphal tip suggesting BafB is localized within endosomes. (G) Staining of cell wall β-1,3-glucan bound to Dectin-1 in Δ*cgnA*^EVOL^; *bafB*^OE-GFP^ shows BafB is localized at the hyphal tip where β-1,3-glucan has yet to be incorporated in the cell wall.

**Figure S6. Overexpression of the *baf* genes alter biofilm morphology in the XY plane of AF293 and CEA10.** (A) Gene expression quantified by qRT-PCR of *hrmA, bafA, bafB,* or *bafC* in AF293 and the respective overexpression strains. n=3 independent biological samples. Error bars indicate standard error of the mean. (B) Gene expression quantified by qRT-PCR of *hrmA, bafA, bafB,* or *bafC* in CEA10 and the respective overexpression strains. n=3 independent biological samples. Error bars indicate standard error of the mean. (C) Representative images from a minimum of three independent biological samples of 24 hour biofilms viewed in the XY dimension for AF293 and the isogenic *baf* overexpression strains. Scale bar is 200 μm. (D) Liquid morphology from 18 hour normoxia-grown cultures. Images are representative of three independent biological samples. (E) Representative images from a minimum of three independent biological samples of 24 hour biofilms viewed in the XY dimension for CEA10 and the isogenic *baf* overexpression strains. Scale bar is 200 μm.

**Figure S7. *Aspergillus niger* encodes a protein similar to *A. fumigatus bafA.*** (A) The protein sequence alignment of *A. fumigatus* BafA (Predicted) with the *A. niger* protein encoded by An0812010. The sequences share 41.03% identity. (B) Introduction of *A. fumigatus bafA* (*AfbafA*) into *A. niger* strain A1144 significantly increases expression. n=3 independent biological replicates. Student’s two-tailed non-parametric t test performed. Error bars indicate standard error around the mean.

**Table S1. Strains used in this study.** The strain name, genotype, and origin for all strains utilized in this study.

**Table S2. Primers used in this study.** Primers used in this study for strain generation and real-time qPCR.

**Video S1**. **Dynamic BafB localization** Two minute time lapse of Δ*cgnA*^EVOL^; *bafB*^OE-GFP^ demonstrating the rapid movement of GFP-labeled BafB (black signal) puncta within the hyphae.

